# The nonstructural protein 5 of coronaviruses antagonizes GSDMD-mediated pyroptosis by cleaving and inactivating its pore-forming p30 fragment

**DOI:** 10.1101/2021.02.23.432418

**Authors:** Fushan Shi, Qian Lv, Tingjun Wang, Jidong Xu, Wei Xu, Yuhua Shi, Xinyu Fu, Tianming Yang, Yang Yang, Lenan Zhuang, Weihuan Fang, Jinyan Gu, Xiaoliang Li

## Abstract

Coronaviruses (CoV) are a family of RNA viruses that typically cause respiratory, enteric and hepatic diseases in animals and humans. Here, we used porcine epidemic diarrhea virus (PEDV) as a model of coronaviruses (CoVs) to illustrate the reciprocal regulation between CoVs infection and pyroptosis. For the first time, we clarified the molecular mechanism of porcine Gasdermin D (pGSDMD)-mediated pyroptosis and demonstrated that amino acids T239 and F240 within pGSDMD-p30 are critical for pyroptosis. Furthermore, 3C-like protease Nsp5 from SARS-CoV-2, MERS-CoV, PDCoV and PEDV can cleave human/porcine GSDMD at the Q193-G194 junction upstream of the caspase-1 cleavage site to produce two fragments which fail to trigger pyroptosis or inhibit viral replication. Thus, we provide clear evidence that coronoviruses may utilize viral Nsp5-GSDMD pathway to help their host cells escaping from pyroptosis, protecting the replication of the virus during the initial period, which suggest an important strategy for coronoviruses infection and sustain.

## Introduction

Coronaviruses (CoVs) are enveloped positive single-strand RNA viruses which belong to the family of *Coronaviridae*. These viruses can cause enteric, respiratory or hepatic diseases in both human and other mamals^1^. According to serological and genotypic characterizations, CoVs are divided into four genera, including *Alphacoronavirs* (α-CoV), *Betacoronavirus* (β-CoV), *Gammacoronavirus* (γ-CoV) and *Deltacoronavirus* (δ-CoV)^2,3^. As a member of the *Alphacoronavirus* genus, porcine epidemic diarrhea virus (PEDV) was first identified in Europe in 1971 which characterized by severe diarrhea, dehydration, vomiting and high mortality in suckling piglets^4^. A highly virulent PEDV reemerged in China in 2010 and spread rapidly in the USA in 2013, causing enormous economic losses to the global pig farming industry^5–7^. The viral genome of PEDV is approximately 28 kb and encodes an accessory protein, two polyproteins and 4 structural proteins. Most of synthesized polyproteins are cleaved by nonstructural protein 5 (Nsp5), a 3C-like protease encoded by ORF1a, and the protease activity of Nsp5 is essential for PEDV’s replication. Nsp5 from different CoVs share highly conserved amino acid sequence which makes Nsp5 as an ideal broad-spectrum antiviral target^8,9^. It has been reported that 3C-like protease of different viruses, including foot-and-mouth disease (FMDV), hepatitis A virus (HAV) and enterovirus 71 (EV71), can antagonize innate immune signaling pathways by disrupting one or more components of the IFN-inducing pathways^10–14^. For coronaviruses, PEDV Nsp5 antagonizes type I IFN signaling by cleaving the nuclear transcription factor kappa B essential modulator (NEMO) at Q231^15^. Porcine deltacoronavirus (PDCoV) Nsp5 cleaves the porcine mRNA-decapping enzyme 1a (pDCP1A) at Q343 to facilitate its replication^16^. A recently published study demonstrates that SARS-CoV-2 Nsp5 can cleave TAB1 and NLRP12 at two distinct cleavage sites^17^. Although many studies have demonstrated the immune evasion strategies of coronaviruses, the molecular mechanism between coronaviruses replication and the innate immune response remains poorly understood.

Pyroptosis is a form of programmed cell death which characterized by cell swelling, pore formation in the plasma, lysis and releases of cytoplasmic contents^18,19^. This type of inflammatory cell death functions as an innate immune effector to antagonize pathogenic microorganisms. Recent studies identified Gasdermin D (GSDMD) as an executioner of pyroptosis, which is cleaved and activated by caspase-1 and caspase-4/5/11^18–20^. Upon caspase-1/4/5/11 cleavage, the N-terminus of GSDMD (GSDMD-p30) can bind to lipids and phosphatidylethanolamine to form pores at 10-20 nm in size and drive to pyroptosis^21–23^. Under the condition of pathogen infection, the pyroptosis helps the host eliminating infected cells and thereby restricts proliferation of intracellular pathogens^24–26^. On the other hand, the 3C-like protease of EV71 virus can facilitate its replication by inhibiting pyroptosis through further cleaving the active GSDMD-p30^27^. However, the relationship between coronaviruses infection and GSDMD-mediated pyroptosis has not been fully illustrated.

In this study, we used PEDV as a model of CoVs to investigate the relationship between CoVs infection and pyroptosis. We found that the pGSDMD-mediated pyroptosis protected host cells against PEDV infection. However, during the early stage of infection, Nsp5 of PEDV directly cleaved pGSDMD at the Q193-G194 junction and produced two inactive fragments, which do not inhibit PEDV replication. Furthermore, we found Nsp5 from other coronaviruses, such as PDCoV, SARS-CoV-2 and MERS-CoV, had activity to cleave both hGSDMD and pGSDMD to inhibit GSDMD-mediated pyroptosis. Therefore, these results demonstrated a previously unknown mechanism of coronaviruses to escape from the pyroptosis of innate immune responses.

## Results

### PEDV infection induces the degradation of pGSDMD

Since GSDMD was reported as a key effector for pyroptosis, many studies had been performed on human and murine GSDMD, but studies focusing on pGSDMD and its function against pathogenic infection were rare. To investigate the role of pGSDMD on pathogenic infection, the amino acid sequence of pGSDMD was predicted and aligned with other GSDMD homologs from human and mouse (Extended Data Fig. 1), and polyclonal antibody against pGSDMD was prepared as previously described (Extended Data Fig. 2)^28,29^.

To determine whether PEDV infection targets pGSDMD, IPEC-J2 cells were infected with PEDV at indicated time points. Cell death was evaluated by LDH release. The results showed that pyroptosis was inhibited at early timepoints post infection (Fig. 1A). Furthermore, PEDV infection induced degradation of pGSDMD in IPEC-J2 cells (Fig. 1B). Similar results were observed in Vero cells transfected with plasmid encoding pGSDMD and infected with PEDV (Fig. 1C and D). In addition, the degradation of pGSDMD induced by PEDV infection increased in an MOI-dependent manner in IPEC-J2 and Vero cells (Fig. 1E and F). These results indicate that pyroptosis and pGSDMD expression are both inhibited by PEDV infection in IPEC-J2 cells and Vero cells.

**Fig. 1.**
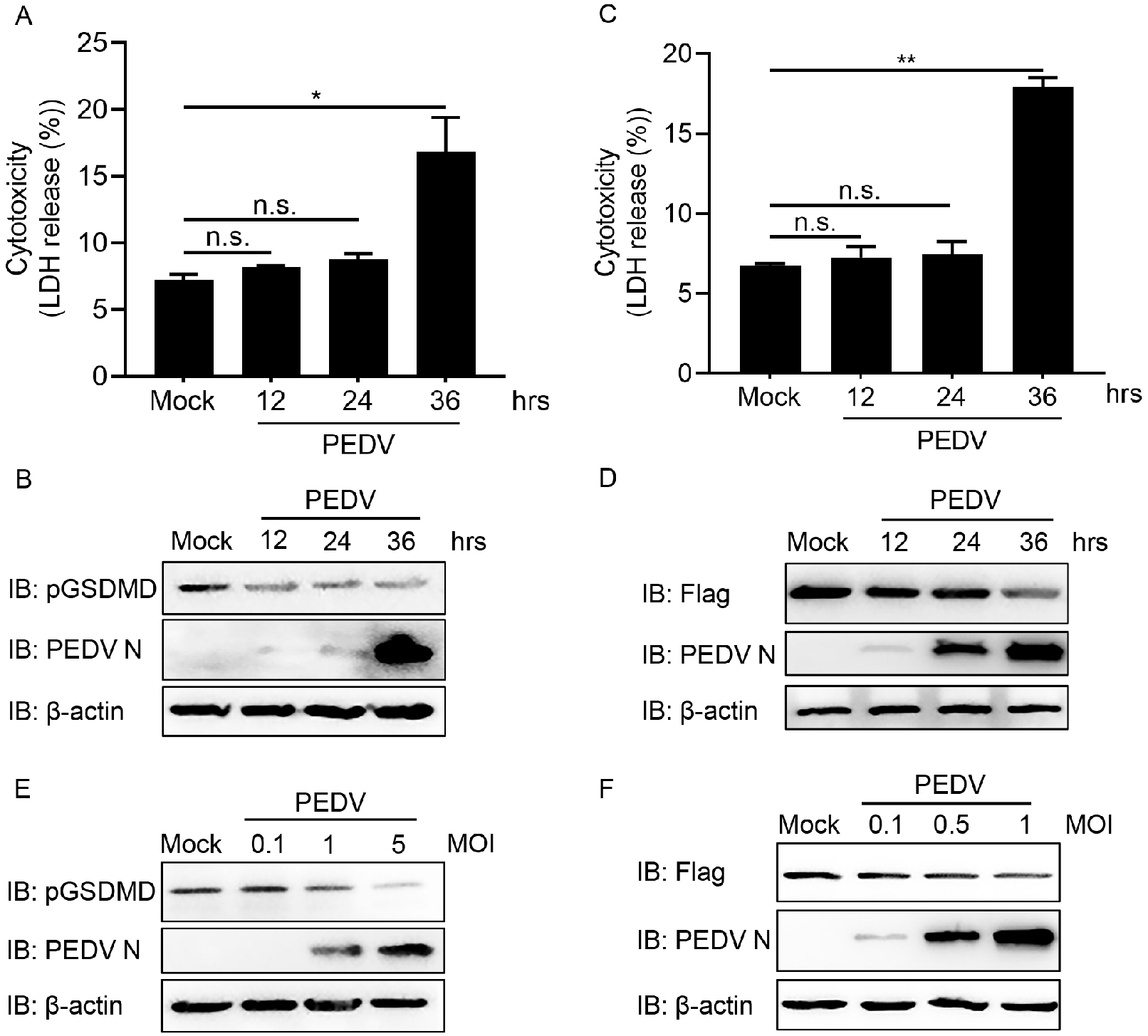
pGSDMD is degraded in PEDV-infected cells. (A and B) IPEC-J2 cells were mock infected or infected with PEDV at an MOI of 1. At the indicated time points, the supernatants were collected and analyzed for LDH level, and cell lysates were processed for Western blotting. n.s., P > 0.05; *, P < 0.05. (C and D) Vero cells were transfected with plasmid encoding p3×Flag-N-GSDMD-FL. At 24 h after transfection, the cells were mock infected or infected with PEDV at an MOI of 0.5. At the indicated time points after infection, the supernatants were collected and analyzed for LDH level, and cell lysates were processed for Western blotting. n.s., P > 0.05; **, P < 0.01. (E) IPEC-J2 cells were mock infected or infected with different doses of PEDV At 24 h after infection, the cells were then processed for Western blotting. (F) Vero cells were transfected with plasmid encoding p3×Flag-N-GSDMD-FL. At 24 h after transfection, the cells were mock infected or infected with different doses of PEDV for another 24 h, and then the cells were processed for Western blotting.

### Caspase-1 cleaves pGSDMD at residue D279-G280 and induces pyroptosis

We next investigate whether pGSDMD could induce pyroptosis. Therefore, porcine caspase-1 (pCaspase-1) and pGSDMD gene were amplified and cloned into vectors to construct recombinant plasmids and then the plasmids were co-transfected into HEK293T cells. As shown in Fig. 2A, co-transfected with plasmids encoding pCaspase-1 and pGSDMD significantly increased the LDH release in HEK293T cells (Fig. 2A). To further confirm the results, the cells were collected and stained with PI, and then analyzed with Fluorescence microscopy and Flow cytometry (Extended Data Fig. 3A and B). Both results showed that co-transfection with plasmids encoding pCaspase-1 and pGSDMD led to increased cell death. These results indicate that co-expression of pGSDMD and pCaspase-1 could induce pyroptosis in HEK293T cells.

**Fig. 2.**
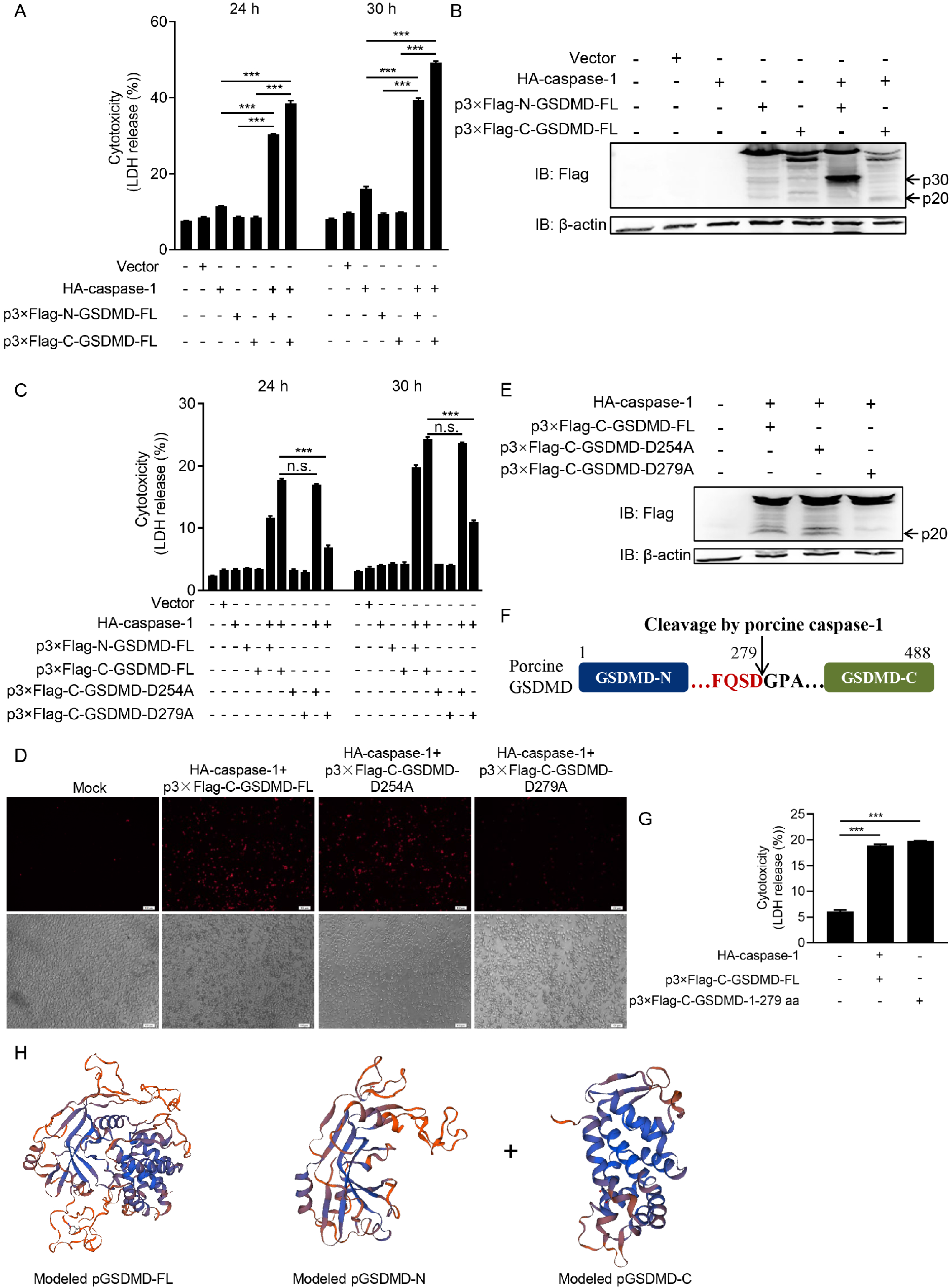
pCaspase-1 mediates pGSDMD cleavage at residue D279. (A and B) HEK293T cells were mock transfected or transfected with plasmids encoding HA-caspase-1 and p3×Flag-N-GSDMD-FL or p3×Flag-C-GSDMD-FL. At the indicated time points after transfection, the supernatants were collected and analyzed for LDH levels (A). At 24 h after transfection, the cells were then processed for Western blotting (B). ***, P < 0.001. (C, D and E) HEK293T cells were transfected with plasmids as shown. At the indicated time points after transfection, the supernatants were collected and analyzed for LDH levels (C). At 24 h after transfection, the cells were staining with PI and analyzed with Fluorescence microscopy (D) or processed for Western blotting (E). ***, P < 0.001; n.s., P > 0.05. (F) Cartoon diagram of porcine GSDMD structure and the cleavage site by caspase-1. (G) HEK293T cells were transfected with plasmids encoding HA-caspase-1 and p3×Flag-C-GSDMD-FL or p3×Flag-C-GSDMD-1-279 aa. At 24 h after transfection, the supernatants were collected and analyzed for LDH levels. ***, P < 0.001. (H) The modeled pGSDMD-FL, pGSDMD-N and pGSDMD-C structure.

The cells were also collected to detect pCaspase-1 mediated cleavage of pGSDMD by Western blotting. As shown in Fig. 2B, cell lysates co-transfected with HA-caspase-1 and p3×Flag-N-GSDMD-FL had a faster-migrating protein band (about 35 kDa) and cell lysates co-transfected with HA-caspase-1 and p3×Flag-C-GSDMD-FL had a smaller protein band (about 25 kDa). These results show that pCaspase-1 could cleave pGSDMD to generate an N-terminal (about 30 kDa) and a C-terminal (about 20 kDa).

Based on the sizes of the cleaved protein bands and the cleavage site preference of caspase-1, the D254-G255 and D279-G280 pairs were tested as the potential cleaved sites for pCaspase-1. Hence, we constructed two mutants in which D was replaced with A. As shown in Fig. 2C and D, wild-type pGSDMD, the D254A mutant and the D279A mutant were co-transfected with pCaspase-1, followed by LDH release assays and PI staining assays. Both results showed that mutation of D279 resulted in significantly decreased pyroptosis, while mutant of D254 showed no significant change, which suggested pCaspase-1 cleaved pGSDMD at residue D279-G280. The Western blotting analysis of the cell lysates was consistent with this result (Fig. 2E), in which the wild-type pGSDMD and the D254A mutant were cleaved by pCaspase-1 and the D279A mutant was resistant to the cleavage. The results indicate that pCaspase-1 cleaves pGSDMD at residue D279-G280 and generated an N-terminal (GSDMD_1-279_) which could induce pyroptosis (Fig. 2F).

To further validate the results, plasmids encoding pGSDMD_1-279_ was constructed and transfected into HEK293T cells. The LDH release assays showed that pGSDMD_1-279_ individually induced pyroptosis (Fig. 2G). Thus, the above results suggest that pGSDMD is cleaved by pCaspase-1 at residue D279-G280 and then generate an N-terminal (p30) which could induce pyroptosis and a C-terminal (p20) (Fig. 2H).

### L295/Y378/A382 are the key sites for pGSDMD autoinhibition

It has been reported that the residues C38/C39 and C191/C192 (human/murine) are essential for oligomerization of GSDMD-N terminal^21,30^. Hence, we next examined whether these key sites also existed on pGSDMD. Based on the multiple-sequence alignment of pGSDMD, hGSDMD and murine GSDMD (mGSDMD) (Extended Data Fig. 1), residues C38 and S191 were tested as the potential key sites for pGSDMD to oligomerize and they were replaced with A to construct point mutants. HEK293T cells were transfected with these point mutants, LDH release results showed that C38A and S191A did not show inhibitory effects on pyroptosis (Fig 3A). However, we further evaluated the oligomerization of pGSDMD-p30 following treatment of NSC and NSA (two specific inhibitors for oligomerization of hGSDMD-p30)^30,31^. As shown in Fig. 3B, the results showed that both NSC and NSA inhibited the pyroptosis induced by porcine/human GSDMD-p30 (Fig. 3B), suggesting that there are residues which impact the oligomerization of pGSDMD-p30 and still required further exploration in the future.

**Fig. 3.**
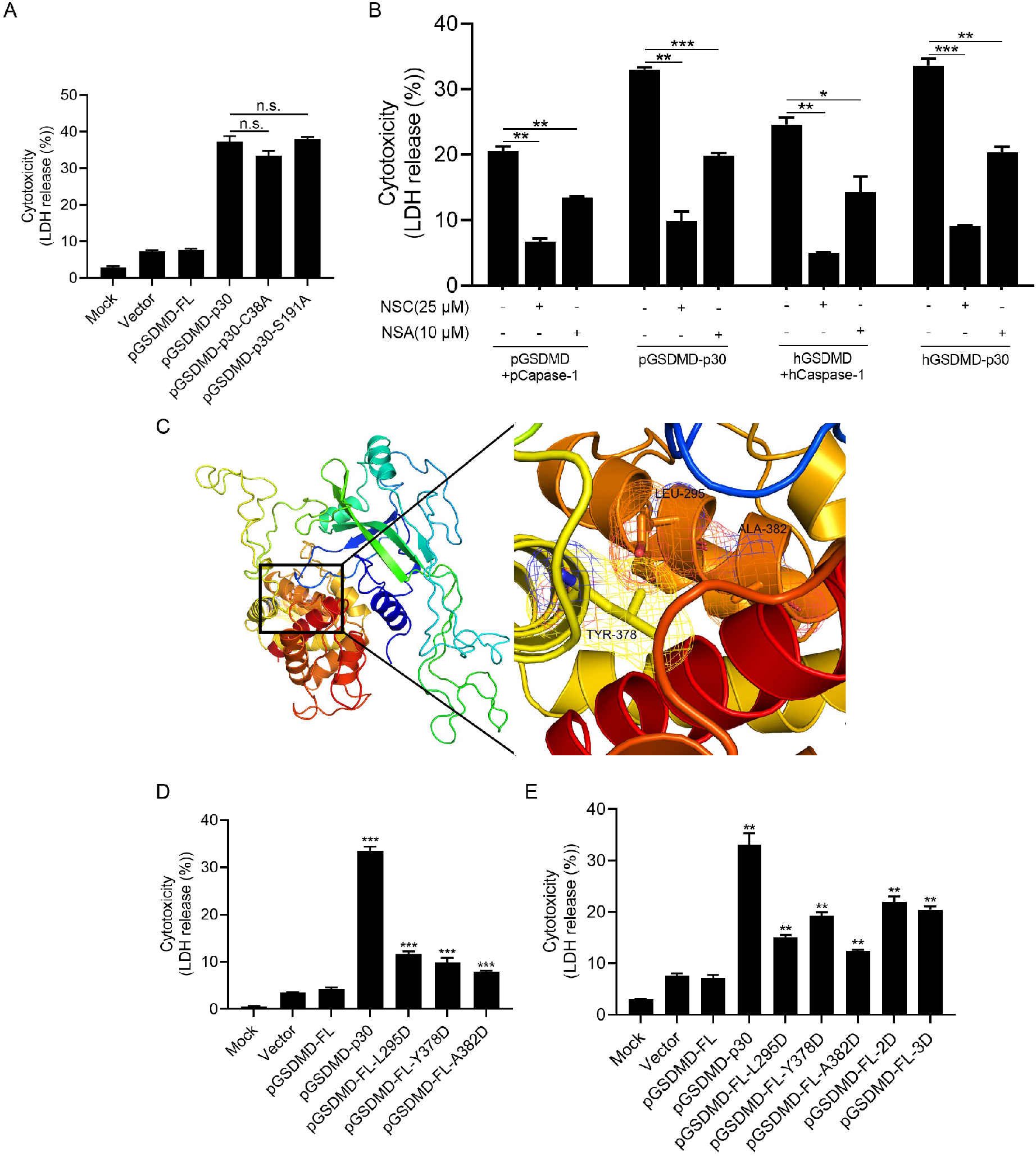
L295/Y378/A382 are the key sites for pGSDMD autoinhibition. (A) HEK293T cells were mock transfected or transfected with plasmids encoding pGSDMD-FL, pGSDMD-p30 and its point mutants. At 24 h after transfection, the supernatants were collected and analyzed for LDH levels. n.s., P > 0.05. (B) HEK293T cells were mock transfected or transfected with plasmids encoding pGSDMD-FL and pCaspase-1, pGSDMD-p30, hGSDMD-FL and hCaspase-1, hGSDMD-p30. Meanwhile, cells were treated with mock, NSC (final concentration of 25 μM) or NSA (final concentration of 10 μM). At 24 h after transfection, the supernatants were collected and analyzed for LDH levels. *, P < 0.05; **, P < 0.01; ***, P < 0.001. (C) The structure of modeled pGSDMD-FL and enlarged view of the boxed area. (D and E) HEK293T cells were mock transfected or transfected with plasmids encoding pGSDMD-p30, pGSDMD-FL and its point mutants. At 24 h after transfection, the supernatants were collected and analyzed for LDH levels. **, P < 0.01; ***, P < 0.001.

Earlier reports have demonstrated that full length of hGSDMD and mGSDMD have an autoinhibitory structure, in which GSDMD-C terminal inhibits the activity of GSDMD-N terminal to induce pyroptosis^23,30^. Based on the multiple-sequence alignment, L295, Y378 and A382 of pGSDMD formed a pocket which associated with GSDMD-N terminal according to the homology modeling results (Fig. 3C). Thus, the three residues were tested as the potential sites in pGSDMD. These residues were separately mutated into D (GSDMD-FL-L295D/GSDMD-FL-Y378D/GSDMD-FL-A382D), L295 and Y373 were simultaneously mutated into D (GSDMD-FL-2D), and three residues were simultaneously mutated into D (GSDMD-FL-3D). The mutants were transfected into HEK293T cells, and results showed that all of the mutants had the activity to induce pyroptosis (Fig. 3D). However, there were no statistic differences between 2D and 3D (Fig. 3E). The aforementioned results suggest that L295, Y378 and A382 are the critical sites for autoinhibitory structure of full length of pGSDMD.

### PEDV Nsp5 associates with and cleaves pGSDMD

To investigate the relationship between PEDV infection and pyroptosis, Vero cells were transfected with plasmids encoding pGSDMD full-length (pGSDMD-FL) and GSDMD N-terminal (pGSDMD-p30) and then infected with PEDV. The LDH release assays results showed that PEDV infection had an inhibition effect on pyroptosis induced by pGSDMD-p30 (Fig. 4A). Meanwhile, the replication of PEDV was also inhibited significantly by pyroptosis induced by pGSDMD-p30 (Fig. 4B). These results suggest that there might be reciprocal regulation between PEDV and the pGSDMD-p30-mediated pyroptosis.

**Fig. 4.**
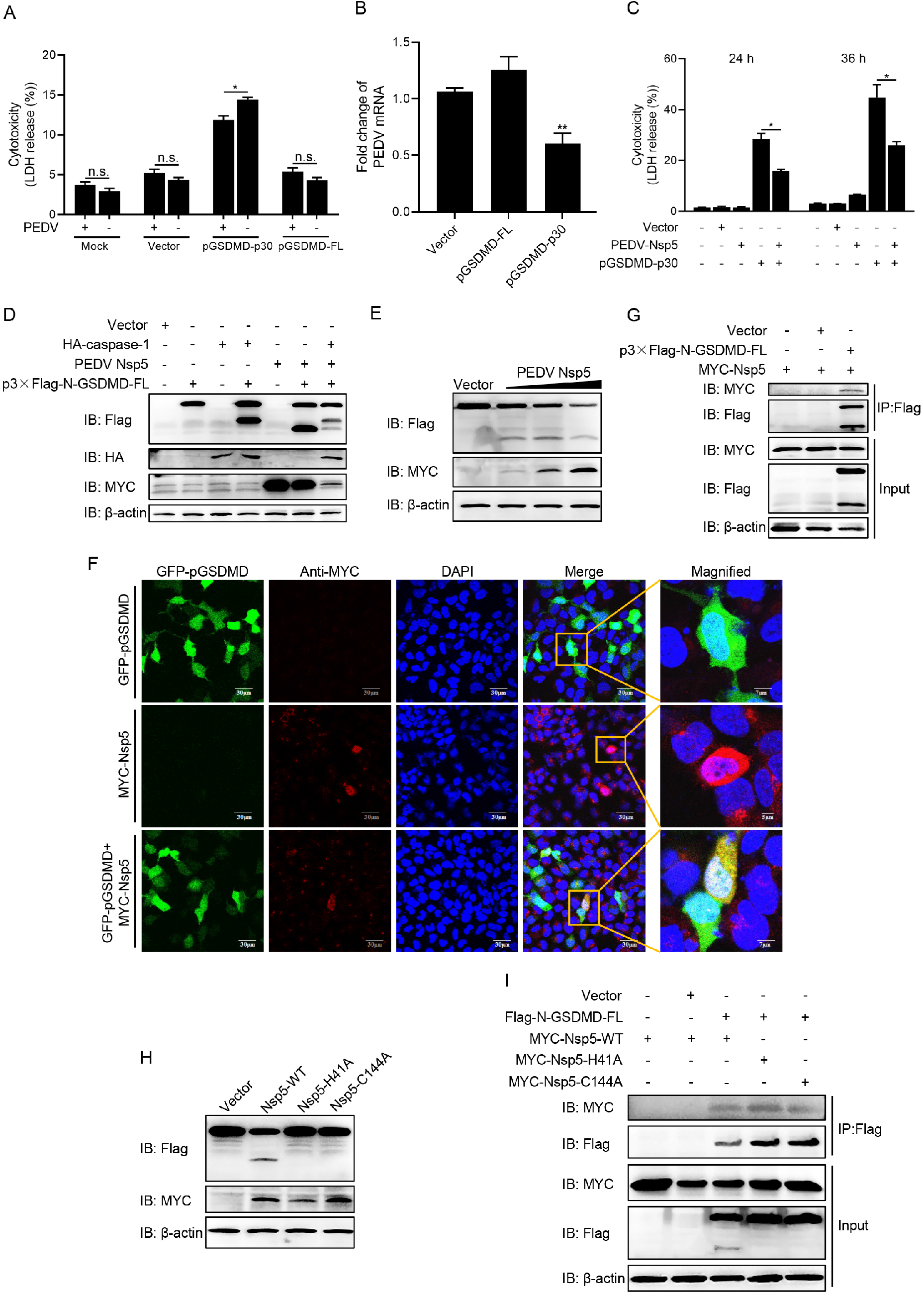
PEDV Nsp5 associates with and cleaves pGSDMD. (A) Vero cells were mock transfected or transfected with plasmids encoding pGSDMD-p30 and pGSDMD-FL. At 4 h after transfection, the cells were mock infected or infected with PEDV at an MOI of 0.1. After 36 h, the supernatants were collected and analyzed for LDH levels. n.s., P > 0.05; *, P < 0.05. (B) Vero cells were transfected with plasmids encoding pGSDMD-p30 and pGSDMD-FL. An empty vector was used as a control. At 24 h after transfection, the cells were infected with PEDV at an MOI of 0.5. After 24 h, total RNA was extracted and the viral RNA levels of PEDV were evaluated by quantitative real-time PCR using SYBR green. Data were expressed as fold change of the PEDV mRNA level relative to that of the control vector. **, P < 0.01. (C) HEK293T cells were transfected with plasmids encoding PEDV-Nsp5 and pGSDMD-p30, or co-transfected with these two plasmids. At 24 h and 36 h after transfection, the supernatants were collected and analyzed for LDH levels. *, P < 0.05. (D) HEK293T cells were transfected with plasmids as shown. At 24 h after transfection, the cells were then processed for Western blotting. (E) HEK293T cells were transfected with plasmids encoding p3×Flag-N-GSDMD-FL and various dose of MYC-Nsp5. After 24 h, cells were lysed for Western blotting. (F) HEK293T cells were transfected with plasmids encoding GFP-pGSDMD and MYC-Nsp5 for 24 h, and then MYC-Nsp5 were labeled with specific primary antibodies and secondary antibodies (red). Cell nuclei were stained with DAPI (blue). The fluorescent signals were observed with confocal immunofluorescence microscopy. HEK293T cells were transfected with plasmids encoding GFP-pGSDMD or MYC-Nsp5 as control. (G) HEK293T cells were transfected with plasmids encoding MYC-Nsp5 and vector or p3×Flag-N-GSDMD-FL for 24 h, followed by CoIP with anti-Flag binding beads and a Western blotting analysis. (H) HEK293T cells were transfected with plasmids encoding p3×Flag-N-GSDMD-FL and wild-type PEDV Nsp5 or its protease-defective mutants (H41A and C144A). After 24 h, cells were lysed for Western blotting. (I) HEK293T cells were transfected with plasmids as shown, followed by CoIP with anti-Flag binding beads and a Western blotting analysis.

Nonstructural protein 5 (Nsp5), the 3C-like protease, which mediates the cleavage of viral polyproteins, has reported be able to cleave a number of host proteins, such as DCP1A and NEMO, to suppress antiviral host responses^10,11,15,16^. Therefore, we speculated that PEDV Nsp5 can cleave pGSDMD to suppress pyroptosis. As shown in Fig. 4C, HEK293T cells were transfected with PEDV Nsp5 or pGSDMD-p30, or co-transfected with these two recombinant plasmids. The supernatants were collected at different time points and tested for LDH release. The results showed that the expression of PEDV Nsp5 inhibited pyroptosis induced by pGSDMD-p30 (Fig. 4C). For further validation, HEK293T cells were transfected with plasmids as indicated in Fig. 4D, and the Western blotting results showed that there was a faster-migrating protein band (about 25 kDa) in samples co-transfected with PEDV Nsp5 and p3 ×Flag-N-GSDMD-FL (Fig. 4D, lane 6), and there were two cleavage protein bands, of 35 kDa (p30) and 25 kDa respectively, in samples co-transfected with HA-caspase-1, PEDV Nsp5 and p3 ×Flag-N-GSDMD-FL (Fig. 4D, lane 7). These results implied that pGSDMD was a target cleaved by PEDV Nsp5. To further confirm the pGSDMD cleavage mediated by PEDV Nsp5, p3×Flag-N-GSDMD-FL was co-transfected with an increasing dose of PEDV Nsp5 into HEK293T cells. Western blotting results showed that pGSDMD cleavage progressively increased in a PEDV Nsp5-dose-dependent manner (Fig. 4E). We next investigated the colocalization of pGSDMD and PEDV Nsp5 with confocal microscopy. As shown in Fig. 4F, HEK293T cells were transfected with plasmids as shown and the protein localization were examined after 24 h. An indirect immunofluorescence assay showed that pGSDMD and Nsp5 colocalized in the cytoplasm (Fig. 4F). The CoIP experiments also demonstrated that PEDV Nsp5 interacted with and cleaved pGSDMD (Fig. 4G).

To further investigate whether PEDV Nsp5 cleaves pGSDMD by means of its protease activity, two Nsp5 mutants, H41A and C144A, which disrupted the protease activity of Nsp5^8,32–34^, were constructed and co-transfected with p3×Flag-N-GSDMD-FL into HEK293T cells. As shown in Fig. 4G, wild-type Nsp5 cleaved pGSDMD successfully, while the two mutants failed to cleave pGSDMD (Fig. 4H). Nevertheless, CoIP experiments showed that the loss of protease activity of Nsp5 did not impact its interaction with pGSDMD (Fig. 4I). Hence, the protease activity of PEDV Nsp5 is essential for pGSDMD cleavage but not interaction.

### PEDV Nsp5 cleaves pGSDMD at residue Q193-G194

We next examined the sequence of pGSDMD for potential PEDV Nsp5 cleavage site. Logo analysis of the cleavage site predicted from the polyprotein cleavage of PEDV Nsp5 was shown in Fig. 5A. Based on the substrate preference of Nsp5 and the sizes of the cleaved bands, the Q193-G194, Q195-G196 and Q197-G198 pairs were tested as the potential cleaved sites^35,36^. Therefore, these three Q residues were replaced with A and these three mutants pGSDMD-Q193A, pGSDMD-Q195A, pGSDMD-Q197A were co-transfected with vector or PEDV Nsp5. As shown in Fig. 5B, Western blotting results showed that pGSDMD-Q193A was resistant to PEDV Nsp5-mediated cleavage, while pGSDMD-Q195A and pGSDMD-Q197A were not (Fig. 5B), suggesting that PEDV Nsp5 cleaves pGSDMD at residue Q193-G194 junction (Fig. 5C).

**Fig. 5.**
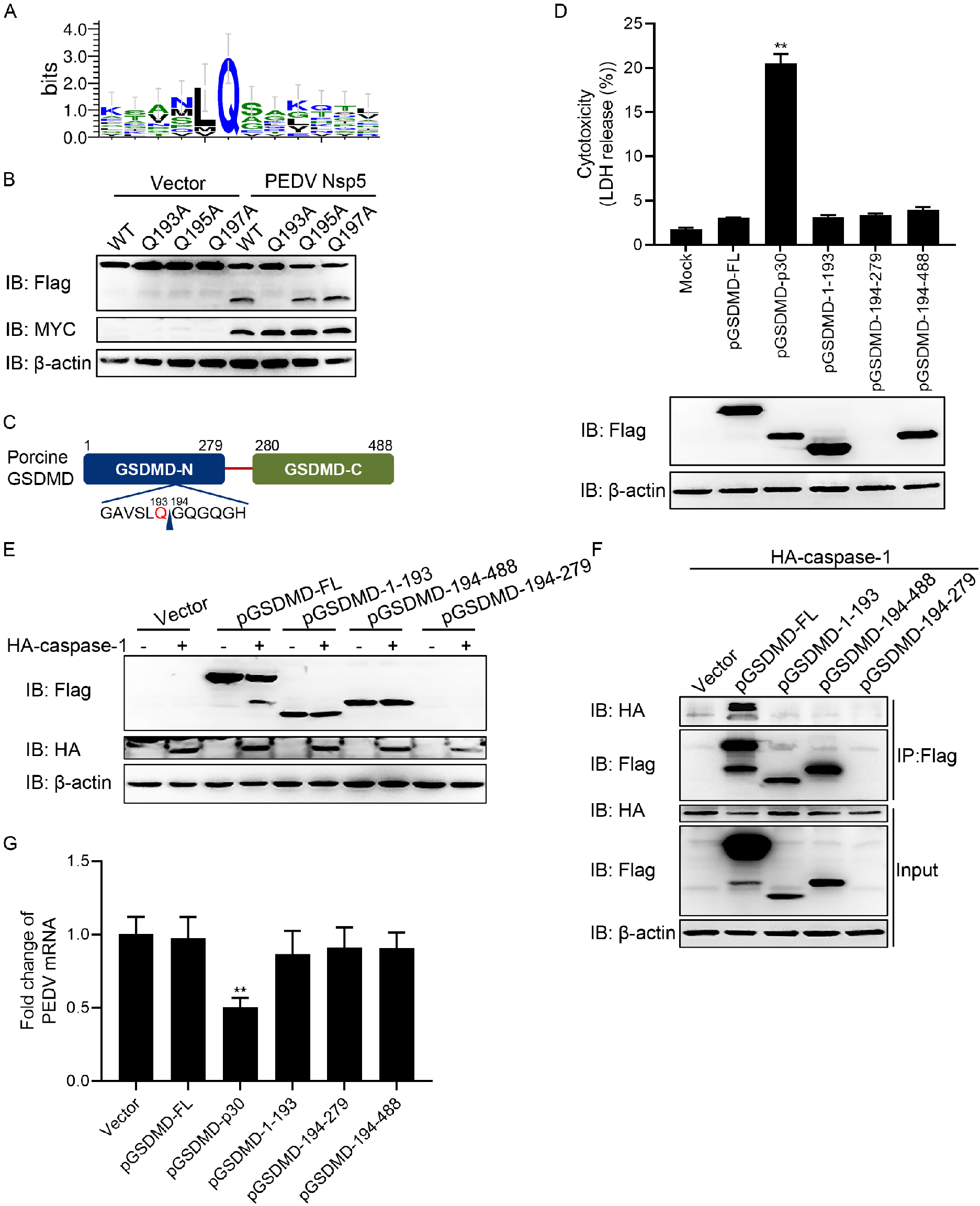
PEDV Nsp5 cleaves pGSDMD at residue Q193. (A) Logo analysis of the predicted cleavage site of PEDV Nsp5. (B) HEK293T cells were transfected with plasmids encoding MYC-Nsp5 and p3×Flag-N-GSDMD-FL or its mutant, including p3×Flag-N-GSDMD-FL-Q193A, p3×Flag-N-GSDMD-FL-Q195A, p3×Flag-N-GSDMD-FL-Q197A. Cells were then lysed after 24 h and evaluated by Western blotting. (C) Cartoon diagram of pGSDMD structure and the cleavage site by PEDV Nsp5. (D) HEK293T cells were mock transfected or transfected with the plasmids encoding pGSDMD-FL, pGSDMD-p30, pGSDMD-1-193, pGSDMD-194-279 and pGSDMD-194-279. After 24 h, the supernatants were collected and analyzed for LDH levels and the cell were then processed for Western blotting. **, P < 0.01. (E) HEK293T cells were transfected with the plasmids as shown. After 24 h, the cells were then processed for Western blotting. (F) HEK293T cells were transfected with plasmids encoding HA-caspase-1 and vector, pGSDMD-FL, pGSDMD-1-193, pGSDMD-194-488, or pGSDMD-194-279, followed by CoIP with anti-Flag binding beads and a Western blotting analysis. (G) Vero cells were transfected with plasmids encoding pGSDMD and its variants as indicated. At 24 h after transfection, cells were infected with PEDV at an MOI of 0.5. After 24 h, total RNA was extracted, and the viral RNA level of PEDV were evaluated by quantitative real-time PCR using SYBR green. **, P < 0.01.

PEDV Nsp5 cleaves pGSDMD to generate pGSDMD1-193 and pGSDMD194-488, and pCaspase-1 cleaves pGSDMD at residue D279. Thus, we next investigated whether these cleaved fragments of pGSDMD_1-193_, pGSDMD_194-279_ and pGSDMD_194-488_ can induce pyroptosis. As shown in Fig. 5D, these three truncated mutants were separately transfected into HEK293T cells and results showed that none of them induced pyroptosis (Fig. 5D). Meanwhile, since the protein band of pGSDMD_194-279_ was too small to be visualized, we subsequently cloned them into EGFP-tagged vectors and then transfected them into HEK293T cells. The results further confirmed that these three truncated mutants cannot induce pyroptosis (Extended Data Fig. 4). Next, we further examined whether pCaspase-1 could associate with and cleave these three truncated mutants. As shown in Fig. 5E and F, HEK293T cells were co-transfected with plasmids as indicated, and the results of CoIP assay and Western blotting assays showed that pCaspase-1 could associate with and cleave full-length of pGSDMD, but had no interaction with pGSDMD_1-193_, pGSDMD_194-279_ and pGSDMD_194-488_.

As described in the preceding text, pyroptosis of cells induced by pGSDMD-p30 had an inhibition effect on PEDV replication. Therefore, we next investigated whether PEDV Nsp5-mediated cleavage of pGSDMD impacts its antiviral activity. Vero cells were transfected with plasmids encoding vector, pGSDMD-FL, pGSDMD-p30, pGSDMD_1-193_, pGSDMD_194-279_ and pGSDMD_194-488_. At 24 h after transfection, cells were infected with PEDV at an MOI of 0.5 for another 24 h, and then total RNA was extracted and the viral RNA level of PEDV were evaluated by quantitative real-time PCR. As shown in Fig. 5G, there was no statistical differences among vector, pGSDMD-FL, pGSDMD_1-193_, pGSDMD_194-279_ and pGSDMD_194-488_, indicating that the antiviral activity of fragments cleaved by Nsp5 were nearly abolished (Fig. 5G). In summary, the above results demonstrate that the antiviral activity of pyroptosis induced by pGSDMD-p30 are impaired by PEDV Nsp5-mediated cleavage, which emphasizes the importance of pGSDMD cleavage on PEDV replication.

### Amino acids T239 and F240 are two key sites for pGSDMD-p30 to induce pyroptosis

It has been shown that pGSDMD_1-279_ (pGSDMD-p30) can induce pyroptosis, while pGSDMD_1-193_ cannot. Based on this, we conjectured that the active motif of pGSDMD to induce pyroptosis located at the amino acids between 193 and 279. Thus, we constructed a series of pGSDMD truncated mutants which encoding pGSDMD_1-254_, pGSDMD_1-244_, pGSDMD_1-234_, pGSDMD_1-224_, and pGSDMD_1-214_, and transfected them into HEK293T cells. As shown in Figure 6A and B, pGSDMD_1-279_ (pGSDMD-p30), pGSDMD_1-254_ and pGSDMD_1-244_ can induce pyroptosis, while pGSDMD_1-234_, pGSDMD_1-224_, pGSDMD_1-214_ cannot (Fig. 6A and B), indicating that the key sites located between amino acids 234 and 244. Hence, the amino acids between 234 and 244 were replaced by D and these point mutants were transfected into HEK293T cells as shown in Fig. 6C. The results showed that all of the point mutants, except T239D and F240D (Fig. 6C), can induce pyroptosis, suggesting that T239 and F240 are the essential sites for pGSDMD-p30 to induce pyroptosis. The results were further proved by PI staining assay (Extended Data Fig. 5). Notably, the point mutant R238D can inhibit the release of LDH but cannot inhibit the intake of PI, suggesting that the mutation of R238 led to smaller pores on cell membrane than wild type pGSDMD-p30.

**Fig. 6.**
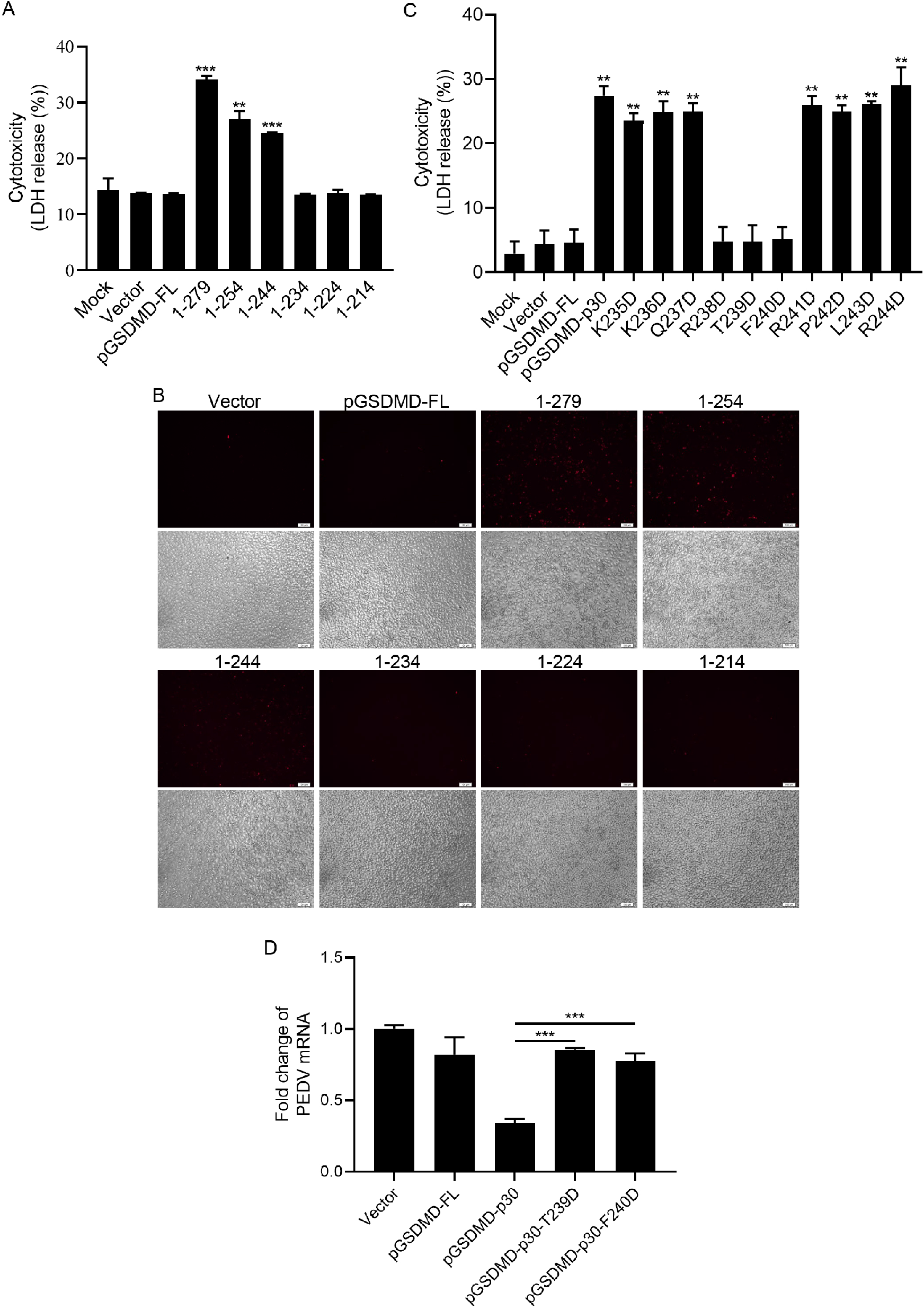
The T239 and F240 amino acids of the N terminus of pGSDMD are necessary for its induced pyroptosis. (A and B) HEK293T cells were transfected with plasmids encoding pGSDMD-FL and its variants. After 24 h, the supernatants were collected and analyzed for LDH levels, and the cells were dyeing with PI. ***, P < 0.001; **, P < 0.01. (C) HEK293T cells were transfected with plasmids encoding pGSDMD-p30 and its point mutants. After 24 h, the supernatants were collected and analyzed for LDH levels. **, P < 0.01. (D) Vero cells were transfected with the plasmids encoding pGSDMD-FL, pGSDMD-p30 and its point mutants (pGSDMD-p30-T239D, pGSDMD-p30-F240D). At 24 h after transfection, cells were infected with PEDV at an MOI of 0.5. After 24 h, total RNA was extracted, and the viral RNA level of PEDV were evaluated by quantitative real-time PCR using SYBR green. ***, P < 0.001.

To further investigate the effects of T239D and F240D on viral replication, the two mutants were transfected into Vero cells along with vector, pGSDMD-FL and pGSDMD-p30, and 24 h after transfection, cells were infected with PEDV at an MOI of 0.5 and then the replication of virus were tested by quantitative real-time PCR. As presented in Fig. 6D, in contrast to pGSDMD-p30, neither T239D nor F240D could inhibit the replication of PEDV (Fig. 6D), further confirming that inhibition of pyroptosis induced by pGSDMD-p30 is essential for PEDV to replicate.

### GSDMD is a common substrate of different coronaviruses Nsp5

Next, we tested whether Nsp5 encoded by other genera of CoVs can cleave GSDMD. Multiple-sequence alignment showed that Nsp5 of PDCoV, SARS-CoV-2 and MERS-CoV were highly similar to PEDV Nsp5 (Extended Data Fig. 6), especially their catalytic domain (Fig. 7A). Thus, the Nsp5 of PDCoV, SARS-CoV-2 and MERS-CoV were cloned into empty vector and co-transfected with pGSDMD, and the Western blotting results showed that all these Nsp5 can cleave pGSDMD (Fig. 7B). To further confirm the results, we respectively constructed point mutants of these Nsp5 which did not show protease activity. As shown in Fig. 7C, PDCoV Nsp5 can cleave pGSDMD while its mutants cannot (Fig. 7C). Likewise, the wild-type Nsp5 of SARS-CoV-2 cleaved both pGSDMD (Fig. 7D) and hGSDMD (Fig. 7E), while its mutants did not. Similar results were observed for cleavage of pGSDMD and hGSDMD by MERS-CoV Nsp5 (Fig. 7F and G). The results above suggest that GSDMD is a common substrate of different genera of coronaviruses Nsp5. To further validate this conclusion, we analyzed the peptides GAVSLQ(193)↓GQGQGH (pGSDMD, arrow represents cleavage site) and Nsp5 of PEDV (Fig. 8A), SARS-CoV-2 (Fig. 8B), MERS-CoV (Fig. 8C) and PDCoV (Fig. 8D), peptides GATCLQ(193)↓GEGQGH (hGSDMD, arrow represents cleavage site) and Nsp5 of SARS-CoV-2 (Fig. 8E) and MERS-CoV (Fig. 8F) by homology modeling^37,38^. As shown in Fig. 8, the residues of both pGSDMD and hGSDMD comfortably fit in the Nsp5 pockets of different CoVs, suggesting a strong interaction between them.

**Fig. 7.**
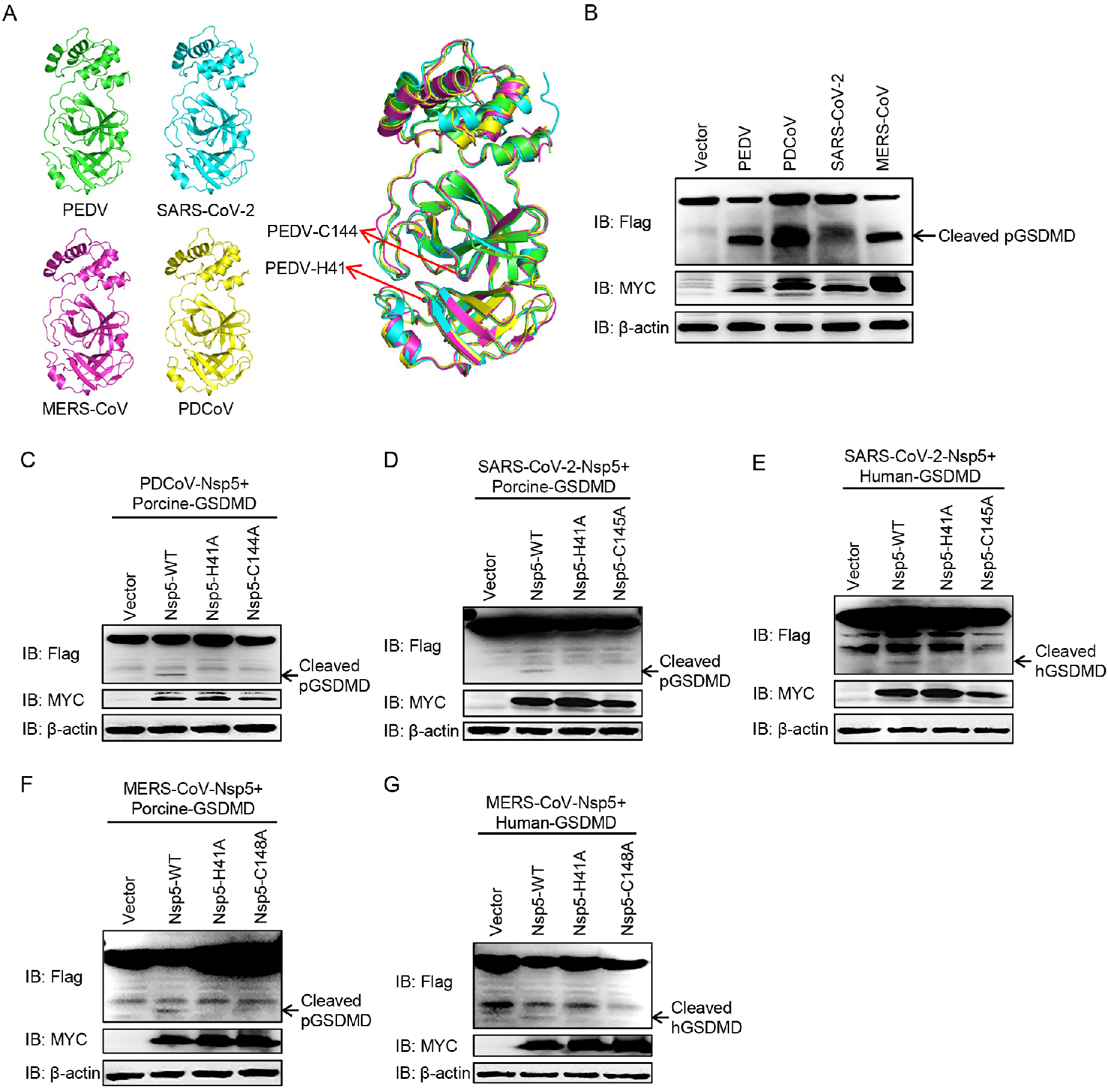
GSDMD is a common target of Nsp5 of different coronaviruses. (A) Structure alignment of CoVs Nsp5. Red arrows indicate conserved enzymatic proteolysis residues His41 and Cys144. The 3D structures were derived from the Protein Data Bank with the following accession numbers: PEDV, 4XFQ; SARS-CoV-2, 7BUY; MERS-CoV, 5WKK; PDCoV, 6JIJ. (B) HEK293T cells were transfected with plasmids encoding p3×Flag-N-GSDMD-FL and Nsp5 encoded by PEDV, PDCoV, SARS-CoV-2, MERS-CoV After 24 h, cells were lysed and detected by Western blotting. (C) HEK293T cells were transfected with plasmids encoding pGSDMD and wild-type PDCoV Nsp5 or its protease-defective mutants (H41A and C144A). After 24 h, cells were lysed for Western blotting. (D and E) HEK293T cells were transfected with plasmids encoding pGSDMD and wild-type SARS-CoV-2 Nsp5 or its protease-defective mutants (H41A and C145A), hGSDMD and wild-type SARS-CoV-2 Nsp5 or its protease-defective mutants (H41A and C145A). After 24 h, cells were lysed for Western blotting. (F and G) HEK293T cells were transfected with plasmids encoding pGSDMD and wild-type MERS-CoV Nsp5 or its protease-defective mutants (H41A and C148A), hGSDMD and wild-type MERS-CoV Nsp5 or its protease-defective mutants (H41A and C148A). After 24 h, cells were lysed for Western blotting.

**Figure 8.**
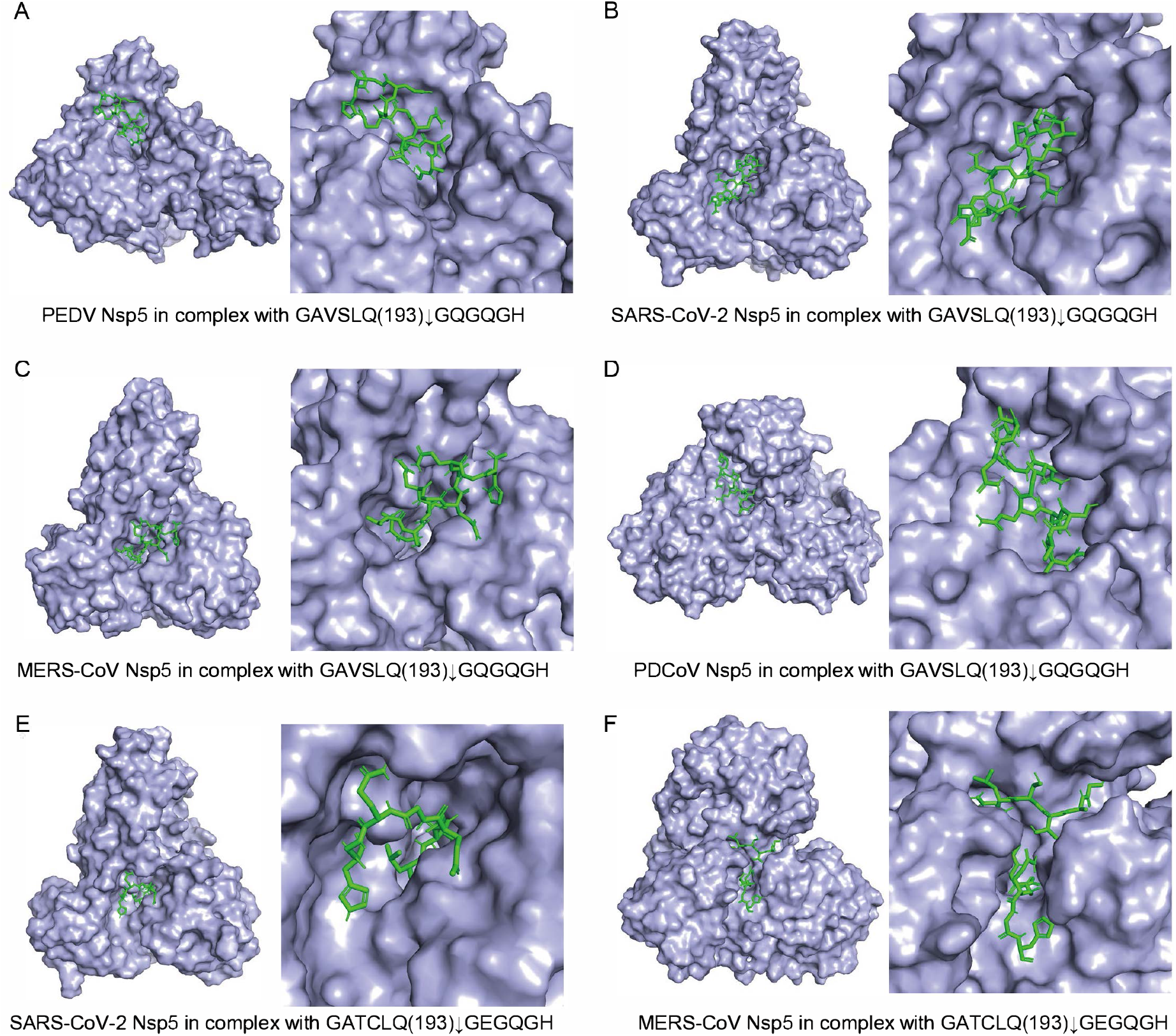
Homology modeling of Nsp5 of different CoVs with the cleaved GSDMD peptide substrate. The molding structure of PEDV Nsp5 (PDB accession number 4XFQ) (A), SARS-CoV-2 (PDB accession number 7BUY) (B and E), MERS-CoV (PDB accession number 5WKK) (C and F), PDCoV (PDV accession number 6JIJ) (D) combined with the cleaved pGSDMD peptide substrate GAVSLQ(193)↓GQGQGH (downward arrows indicates cleavage sites) (A, B, C and D) and hGSDMD peptide substrate GATCLQ(193)↓GEGQGH (downward arrows indicates cleavage sites) (E and F) were analyzed using PyMOL software.

## Discussion

Although considerable progress has been made in CoVs research, knowledge gaps still exist with respect to the host innate immune responses against CoVs infection. Here, we used PEDV as a model of CoVs to illustrate the relationship between CoVs infection and pyroptosis (Fig. 9). We demonstrated that the pGSDMD plays a protective role against PEDV infection. At early time points after PEDV infection, pGSDMD was cleaved by Nsp5 to produce two inactive fragments which failed to trigger pyroptosis and inhibited PEDV replication. The pGSDMD-p30 could inhibit PEDV replication, and the amino acids T239 and F240 determined its inhibitory effect. Furthermore, Nsp5 of other genera can also cleave hGSDMD and pGSDMD to produce inactive fragments. Thus, our results demonstrated that GSDMD may be an appealing target for the design of anti-coronavirus therapies.

**Figure 9.**
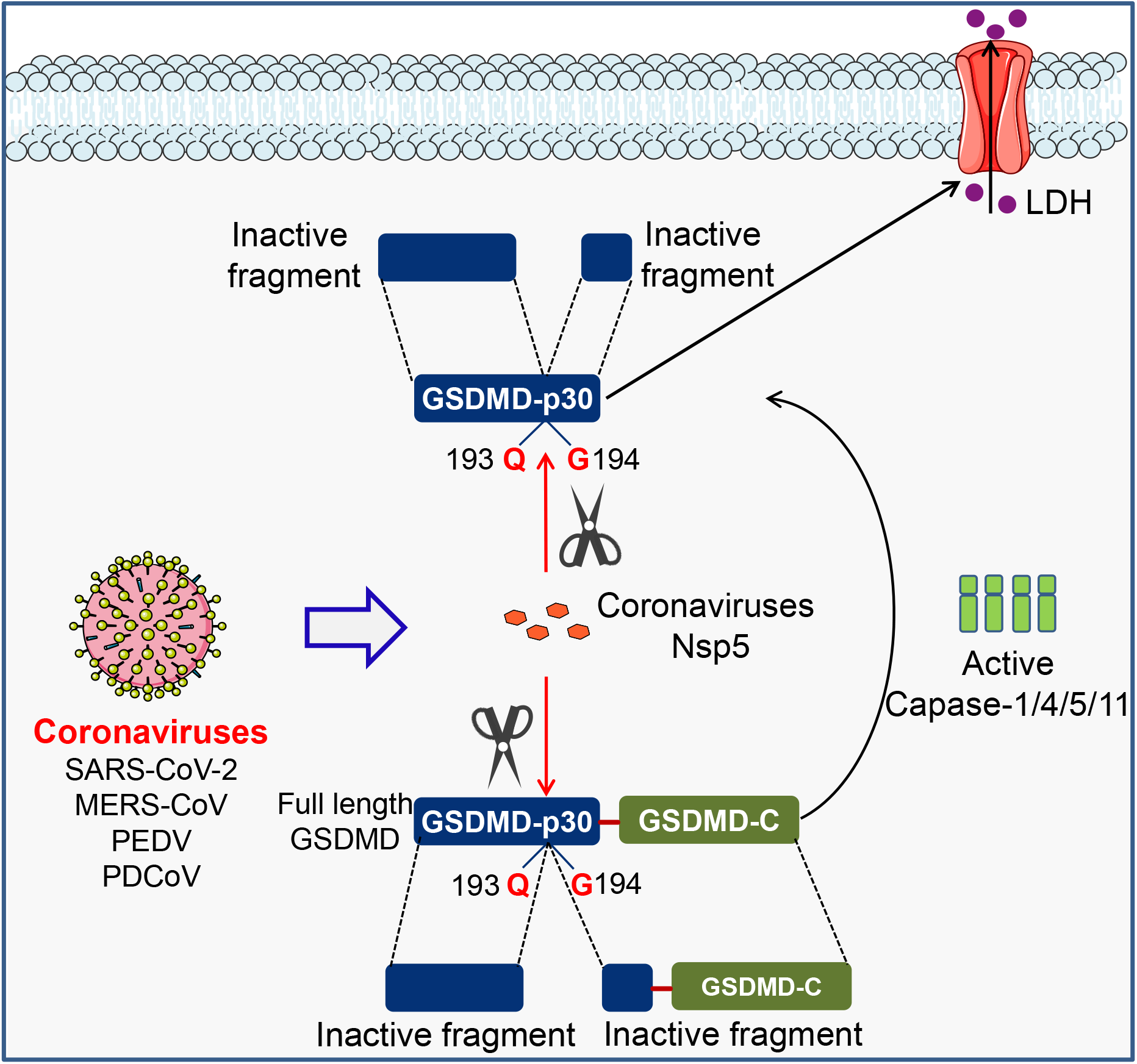
Diagram of CoVs antagonize GSDMD-mediated pyroptosis.

Recent studies have identified that human/murine GSDMD is a direct substrate of caspase-1/4/5/11 and serves as the executioner for pyroptosis. However, the amino acids sequence and molecular characterization of pGSDMD have not been illustrated. In order to investigate the role of pGSDMD-mediated pyroptosis in PEDV infection, we first clarified the molecular characterization of pGSDMD. Porcine GSDMD has 488 aa and can be cleaved by porcine caspase-1 at D279 to produce GSDMD-NT (p30, 1-279 aa), which lead to pyroptosis. Site-directed mutagenesis studies revealed that C38/C39 and C191/C192 (human/murine) mutations impaired hGSDMD-p30/mGSDMD-p30 oligomerization, which is critical for hGSDMD-p30/mGSDMD-p30 during pore formation^22,30^. However, our results indicated that mutation of porcine C38 or S191 (corresponding to human C38 and C191) had no effect on p30-induced pyroptosis. Interestingly, inhibitors of hGSDMD-p30 oligomerization could also abrogate pGSDMD-p30-induced pyroptosis. The results suggest that other critical site(s) determine(s) pGSDMD-p30 oligomerization. Furthermore, our results demonstrated that T239 and F240 within pGSDMD-p30 are critical for inducing pyroptosis.

Generally, the 3C-like protease of CoVs is critical for viral replication by cleaving polyprotein precursors to produce mature nonstructural proteins. However, the 3C-like protease has also acquired mechanisms to evade host innate immune responses. It is reported that CoVs Nsp5 can antagonize innate immune signaling pathways by disrupting one or more components of the IFN-inducing pathways^10–12^. Our present study first demonstrated that CoVs Nsp5 can subvert innate immune responses by cleaving and inactivating GSDMD. Thus, GSDMD represents a novel target of CoVs Nsp5. PEDV Nsp5 not only interacted with and cleaved full length of pGSDMD, but also abrogated pGSDMD-p30-induced pyroptosis by cleaving the p30 fragment. Conversely, protease-dead mutants of the four CoVs Nsp5 were unable to cleave human/porcine GSDMD. Thus, these results suggest a reciprocal regulation between CoVs Nsp5 and pyroptosis.

It is noteworthy that CoVs Nsp5 cleaves human/porcine GSDMD at the Q193-G194 junction. Our results suggest that amino acids T239 and F240 within pGSDMD-p30 are critical for pyroptosis. These two sites within pGSDMD-p30 determine CoVs replication. Upon cleavage by CoVs Nsp5, the truncated N-terminal fragment without T239 and F240 sites failed to induce pyroptosis or inhibit viral replication. Interestingly, a newly published study demonstrated that Zika virus (ZIKV) protease directly cleaved the hGSDMD into N-terminal fragment (1-249), which contains T239 and F240. ZIKV NS2B3 protease cleaves hGSDMD at residue R249 to produce hGSDMD1-249 fragment, which lead to pyroptosis in a caspase-independent manner^39^. Consistent with this, a previous study demonstrated that NS5 protein of ZIKA could directly interact with NLRP3 protein and facilitate NLRP3 inflammasome activation^40^, which is an upstream event for hGSDMD-p30-mediated pyroptosis. Therefore, viruses use different strategies to evade host immune responses and facilitate its replication.

In summary, we used PEDV as a model of coronaviruses to illustrate the reciprocal regulation between CoVs infection and pyroptosis. For the first time, we clarified the molecular mechanism of pGSDMD-mediated pyroptosis and demonstrated that amino acids T239 and F240 within pGSDMD-p30 are critical for pyroptosis. Furthermore, 3C-like protease Nsp5 from SARS-CoV-2, MERS-CoV, PDCoV and PEDV can cleave human/porcine GSDMD at the Q193-G194 junction upstream of the caspase-1 cleavage site to produce two fragments which fail to trigger pyroptosis or inhibit viral replication. Thus, we provide clear evidence that the coronoviruses might utilize its Nsp5 to escape the host pyroptotic cell death in favor of its replication during the initial period, an important strategy for their sustaining infections. Further work is needed to investigate the role of GSDMD in viral pathogenesis.

## Methods

### Plasmids and antibodies

The pGSDMD gene was amplified from cDNA of IPEC-J2 cells by PCR and then cloned into p3×Flag-CMV-7.1 vector, p3×Flag-CMV-14 vector and pEGFP-C1 vector, respectively. Nsp5 of PEDV was amplified from cDNA of PEDV and cloned into PRK5-MYC vector. The truncation mutants and point mutants were generated by PCR of the corresponding plasmids. All of the plasmids were constructed by homologous recombination using ClonExpress II One Step Cloning Kit (Vazyme, C112), and all of the point mutants were constructed using Mut Express II Fast Mutagenesis Kit V2 (Vazyme, C214) and DpnI endonuclease (NEB, R0176S). Primers used for plasmids construction were listed in Supplementary table 1 and table 2. All plasmids were verified by sequencing.

Anti-Flag antibody (F1804), anti-MYC antibody (C3956) and anti-GSDMD antibody (G7422) were purchased from Sigma. Anti-HA antibody (3724) was purchased from Cell Signaling Technology. Anti-β-actin antibody (A01010) was purchased from Abbkine. Anti-GSDMDC1 antibody (sc-393581) was purchased from Santa Cruz. The anti-PEDV N monoclonal antibody and the anti-GSDMD polyclonal antibody were prepared in our laboratory as previously described^28,29^. Necrosulfonamide (S8251) and Disulfiram (S1680) were purchased from Selleck.

### Cells and virus

African green monkey kidney cells (Vero cells) and Human embryonic kidney 293T cells (HEK293T) were cultured in DMEM (Hyclone, SH30243.01) containing 10% fetal bovine serum (FBS) (Corille, C1015-05) and 5% Penicillin-Streptomycin Solution (Hyclone, SV30010). IPEC-J2 cells were maintained in DMEM/F12 (Hyclone, SH30023.01) supplemented with 10% FBS and 5% Penicillin-Streptomycin Solution. Cells were incubated at 37°C with 5% CO2. When cells seeded in cell culture plates grown to approximately 60%, they were transfected with plasmids using VigoFect (Vigorous Biotechnology, T001) or Lipo8000 Transfection Reagent (Beyotime, C0533) according to the manufacturer’s instructions.

The PEDV strain ZJ15XS0101 (GenBank accession KX550281) was isolated and stored in our laboratory^41^. Vero cells and IPEC-J2 cells grown to approximately 80%-90% in cell culture plates were infected with PEDV at different dose with 4 μg/mL trypsin.

### Cytotoxicity assays

Cell death were measured using CytoTox 96 Non-Radioactive Cytotoxicity Assay kit (Promega, G1780) according to the lactate dehydrogenase (LDH) released into medium.

### Western blotting

Cells were lysed with RIPA Lysis Buffer (Beyotime, P0013B) containing PMSF (Beyotime, ST506) for 10 min at 4□ and then denatured in 5 × SDS-PAGE loading buffer (Solarbio, P1040) for 10 min. After harvest, the cell lysate of equal amount was loaded on 8%-12% SDS-PAGE gels (Fdbio science) and electrophoresed, and then transferred to polyvinylidene difluoride membranes (BIO-RAD, 1620177). The membranes were blocked with QuickBlock Blocking Buffer for Western blotting (Beyotime, P0252) for 1 h and then incubated with primary antibodies diluted with QuickBlock Primary Antibody Dilution Buffer for Western blotting (Beyotime, P0256) at 4□ overnight. Membranes were washed with TBST for 10 min (3 times) and then incubated with secondary antibodies diluted in TBST for 1h at room temperature. After washed 3 times with TBST (10 min each), the chemiluminescent signals were analyzed with Clinx imaging system (Clinx Science Instruments).

### Propidium iodide assays

HEK293T cells were seeded in 24-well plates and transfected with indicated plasmids for 24 h. After that, the medium was collected for cytotoxicity assays. The cells were washed for 3 times gently and stained with propidium iodide (BD Bioscience, 556463) and then analyzed with fluorescence microscopy.

### Flow cytometry assays

Cells were harvested using trypsin and washed with PBS 3 times gently and then stained with propidium iodide (BD Bioscience, 556547) according to the manufacturer’s instructions. The cells were analyzed with Flow cytometer (Becton Dickinson, FACSVerse).

### RNA extraction and RT-qPCR

To collect the RNA of PEDV, the medium was discarded and total RNA of the cells and virus was extracted with RNA-easy Isolation Reagent (Vazyme, R701-01). After measuring the concentration of extracted RNA, reverse transcription was conducted with HiScript III 1st Strand cDNA Synthesis Kit (+gDNA wiper) (Vazyme, P312-02) according to the manufacturer’s instructions. Afterwards, cDNA samples were analyzed by qPCR using ChamQ Universal SYBR qPCR Master Mix (Vazyme, P711-01). Primers used for RT-qPCR were listed in Supplementary table 3.

### CoIP assay

HEK293T cells seeded in 6-well plates were transfected with the specific plasmids for 24 h and then cells were lysed with Cell lysis buffer for Western and IP (Beyotime, P0013) containing PMSF (Beyotime, ST506) or Protease inhibitor cocktail for general use (Beyotime, P1005) for 30 min on the ice. Afterwards, the lysates were centrifuged at 4 □ and the supernatants were incubated with anti-Flag binding beads (Sigma, M8823) at 4 □ overnight. The binding beads were then washed with TBS 5 times and then denatured in 1 × SDS-PAGE loading buffer for 10 min. Finally, the supernatants were analyzed with Western blotting.

### Confocal immunofluorescence assay

HEK293T cells were seeded in 24-well plates on coverslips and after cultured overnight indicated plasmids were transfected. At 24 h after transfection, cells were washed 3 times with cold PBS and then fixed with Immunol Staining Fix Solution (Beyotime, P0098) at room temperature for 30 min, following with washing 3 times with PBS (2 min each). Then the cells were permeabilized with Immunostaining Permeabilization Solution with Saponin (Beyotime, P0095) at room temperature for 20 min and washed with PBS 3 times (5 min each). After that, cells were blocked with QuickBlock Blocking Buffer for Immunol Staining (Beyotime, P0260) at room temperature for 60 min, and then incubated with primary antibody (anti-MYC, Sigma, C3956) at 4 □ overnight. After washing 3 times with PBS (3 min each), the cells were incubated with the secondary antibody (Goat Anti-Rabbit IgG Alexa Fluor 568, Abcam, ab175471) at 37 □ for 60 min in the dark and then they were washed 3 times with PBS (5 min each). Nuclei were stained with DAPI (Beyotime, C1002) for 5 min and then cells were washed 4 times with PBS (5 min each). The cells were then analyzed with a Laser confocal microscopy (Olympus, IX81-FV1000).

### Sequence alignment

We collected amino acid sequence of pGSDMD and other GSDMD homologs from human (GenBank accession NP_001159709.1) and mouse (GenBank accession 6N9N_A), and the amino acid sequence of Nsp5 of PEDV and Nsp5 of PDCoV (GenBank accession AKQ63081.1), SARS-CoV-2 (GenBank accession NC_045512) and MERS-CoV (GenBank accession NC_038294). SnapGene software were used to perform the multiple-sequence alignment.

### Statistical analysis

All experiments were repeated three times or more. Data are presented as mean ± SD and analyzed by the two-tailed Student’s *t* test or one-way ANOVA followed by Tukey’s multiple comparisons test by Prism software (GraphPad). The differences were considered significant when*p* < 0.05 (*), *p* < 0.01 (**), and*p* < 0.001 (***).

## Acknowledgements

This work was financially supported by grants from the National Natural Science Foundation of China (32072817), the National Key Research & Development Program of China (2016YFD0500102), the Zhejiang Provincial Key R&D Program of China (2021C02049), the Scientific Research Fund of Zhejiang Provincial Education Department (Y202045613), the Zhejiang Provincial Natural Science Foundation of China (LY18C180001, LY21C180001), and the Fundamental Research Funds for the Central Universities of China (2020XZZX002-20).

We thank Dr. Ying Shan in the Shared Experimental Platform for Core Instruments, College of Animal Sciences, Zhejiang University for assistance with analysis of laser confocal microscopy imaging.

## Author contributions

F. Shi, W. Fang, X. Li and Q. Lv conceived the overall scope of the project. F. Shi and Q. Lv designed and performed the majority of the experiments. J. Gu and T. Wang performed the viral culture and the preparation of polyclonal antibodies; J. X helped with quantitative real-time PCR; W. Xu assisted Q. Lv in Flow cytometry; Y Shi, X. Fu and T. Yang assisted Q. Lv in plasmid construction and confocal immunofluorescence assay; Y. Yang helped with statistical analysis; L. Zhuang assisted F. Shi and Q. Lv wrote the manuscript. All authors discussed the results and reviewed the manuscript.

## Competing interests

The authors declare no competing interests.

---

**Extended Data Figure 1** Alignment of the amino acid sequence of pGSDMD and other GSDMD homologs from human (GenBank accession NP_001159709.1) and mouse (GenBank accession 6N9N_A).

**Extended Data Figure 2** HEK293T cells were mock transfected or transfected with plasmids encoding p3×Flag-N-GSDMD-FL. At 24 h after transfection, cell lysates were analyzed by Western blotting with antibodies for Flag, β-actin and the polyclonal antibody directed against pGSDMD prepared in our laboratory.

**Extended Data Figure 3** HEK293T cells were mock transfected or transfected with plasmids as shown. At 24 h after transfection, the cells were processed and staining with PI, and then analyzed with Fluorescence microscopy (A) and Flow cytometry (B).

**Extended Data Figure 4** HEK293T cells were mock transfected or transfected with the plasmids encoding EGFP-GSDMD-FL, EGFP-GSDMD-p30, EGFP-GSDMD-1-193, EGFP-GSDMD-194-279, EGFP-GSDMD-194-488. After 48 h, the supernatants were collected and analyzed for LDH levels, and the cells were analyzed with Fluorescence microscopy. *, P < 0.05.

**Extended Data Figure 5** HEK293T cells were transfected with plasmids encoding pGSDMD-p30 and its point mutants. After 24 h, the cells were dyeing with PI and analyzed with Fluorescence microscopy.

**Extended Data Figure 6** Alignment of the amino acid sequence of Nsp5 of PEDV with Nsp5 of PDCoV (GenBank accession AKQ63081.1), SARS-CoV-2 (GenBank accession NC_045512) and MERS-CoV (GenBank accession NC_038294).

**Supplementary Tables** Primers used in this study.

## Notes

### Competing Interest Statement

The authors have declared no competing interest.

