## Extended Data Figure 1 for "The nonstructural protein 5 of coronaviruses antagonizes GSDMD-mediated pyroptosis by cleaving and inactivating its pore-forming p30 fragment"

Extended Data Fig. 1

```
        1        10        20        30        40        50
Porcine MASAFERVVKSVVRELDHG.RELTPVKSLQTSDRFQPYCLLGRKPSSSWFWRPRYTCVDL
Human   MGSAFERVVRRVVQELDHG.GEFIPVTSLQSSTGFQPYCLVVRKPSSSWFWKPRYKCVNL
Mouse   MPSAFEKVVKNVIKEVSGSRGDLIPVDSLRNSTSFRPYCLLNRKFSSSRFWKPRYSCVNL

        60        70        80        90        100       110
Porcine SIWDILEPSAPEPAVERGGPFYFHDTMDGQLQSQVELAAPGQGKFSGGAAVSGSSSASMN
Human   SIKDILEPDAAEPDVQRGRSFHFYDAMDGQIQGSVELAAPGQAKIAGGAAVSDSSSTSMN
Mouse   SIKDILEPSAPEPEPECFGSFKVSDVVDGNIQGRVMLSGMGEGKISGGAAVSDSSSASMN

        120       130       140       150       160       170
Porcine VCTLRVAPNTWDAMHLERRLRQPEHKVLQQLRSRGNDVFVVTEVLQTQKEVEVTRTHKQE
Human   VYSLSVDPNTWQTLLHERHLRQPEHKVLQQLRSRGDNVYVVTEVLQTQKEVEVTRTHKRE
Mouse   VCILRVTQKTWETMQHERHLQQPENKILQQLRSRGDDLFVVTEVLQTKEEVQITEVHSQE

        180       190       200       210       220       230
Porcine GSGQFALPGAVSLQGQGQGHLSRKKTVTIPSGSVIAFRVAQLVIGSDWDILLFPDKKQRT
Human   GSGRFSLPGATCLQGEGQGHLSQKKTVTIPSGSTLAFRVAQLVIDSDLDVLLFPDKKQRT
Mouse   GSGQFTLPGALCLKGEGKGHQSRKKMVTIPAGSILAFRVAQLLIGSKWDILLVSDEKQRT

        240       250       260       270       280       290
Porcine FRPLREGHSASHGADGQPPQFSRLVSGMSFPSEHLKFQSDG.PAEDQLVTTEDFQGLQAE
Human   FQPPATGHKRST.SEGAWPQ...LPSGLSMMRCLHNFLTDGVPAEGAF..TEDFQGLRAE
Mouse   FEPSSGDRK....AVGQRHHGLNVLAALCSIGKQLSLLSDG.IDEEELIEAADFQGLYAE

        300       310       320       330       340       350
Porcine VDAWAAGLEALSREPCRQLLGGLGQVLQDEPALQALEDSLEQGLC.GGLVEPQDGPAGAM
Human   VETISKELELLDRELCQLLLEGLEGVLRDQLALRALEEALEQGQS.LGPVEPLDGPAGAV
Mouse   VKACSSELESLEMELRQQILVNIGKILQDQPSMEALEASLGQGLCSGGQVEPLDGPAGCI

        360       370       380       390       400       410
Porcine LECLVLACRRLEREVAGPIFYLVQALAVLNETQHVLLAKVLETGALSGALTLVGSLLEQS
Human   LECLVLSSGMLVPELAIPVVYLLGALTMLSETQHKLLAEALESQTLLGPLELVGSLLEQS
Mouse   LECLVLDSGELVPELAAPIFYLLGALAVLSETQQQLLAKALETTVLSKQLELVKHVLEQS

        420       430       440       450       460       470
Porcine APWQERRAVSLPPTLLGSSWGSEAPIWALLEECGLEPQVGTAQVCWEPEAQSCTCALYAC
Human   APWQERSTMSLPPGLLGNSWGEGAPAWVLLDECGLELGEDTPHVCWEPQAQGRMCALYAS
Mouse   TPWQEQSSVSLPTVLLGDCWDEKNPTWVLLEECGLRLQVESPQVHWEPTSLIPTSALYAS

        480
Porcine LALLLRLSQ.LC
Human   LALLSGLSQEPH
Mouse   LFLLSSLGQKPC
```
