## Supplementary figures and images for "The nonstructural protein 5 of coronaviruses antagonizes GSDMD-mediated pyroptosis by cleaving and inactivating its pore-forming p30 fragment"

### Extended Data Figure 2

Extended Data Fig. 2

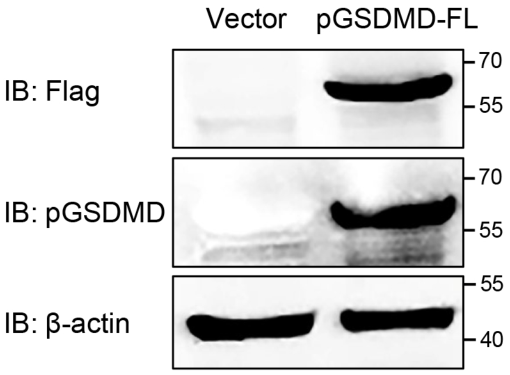

### Extended Data Figure 3

Extended Data Fig. 3

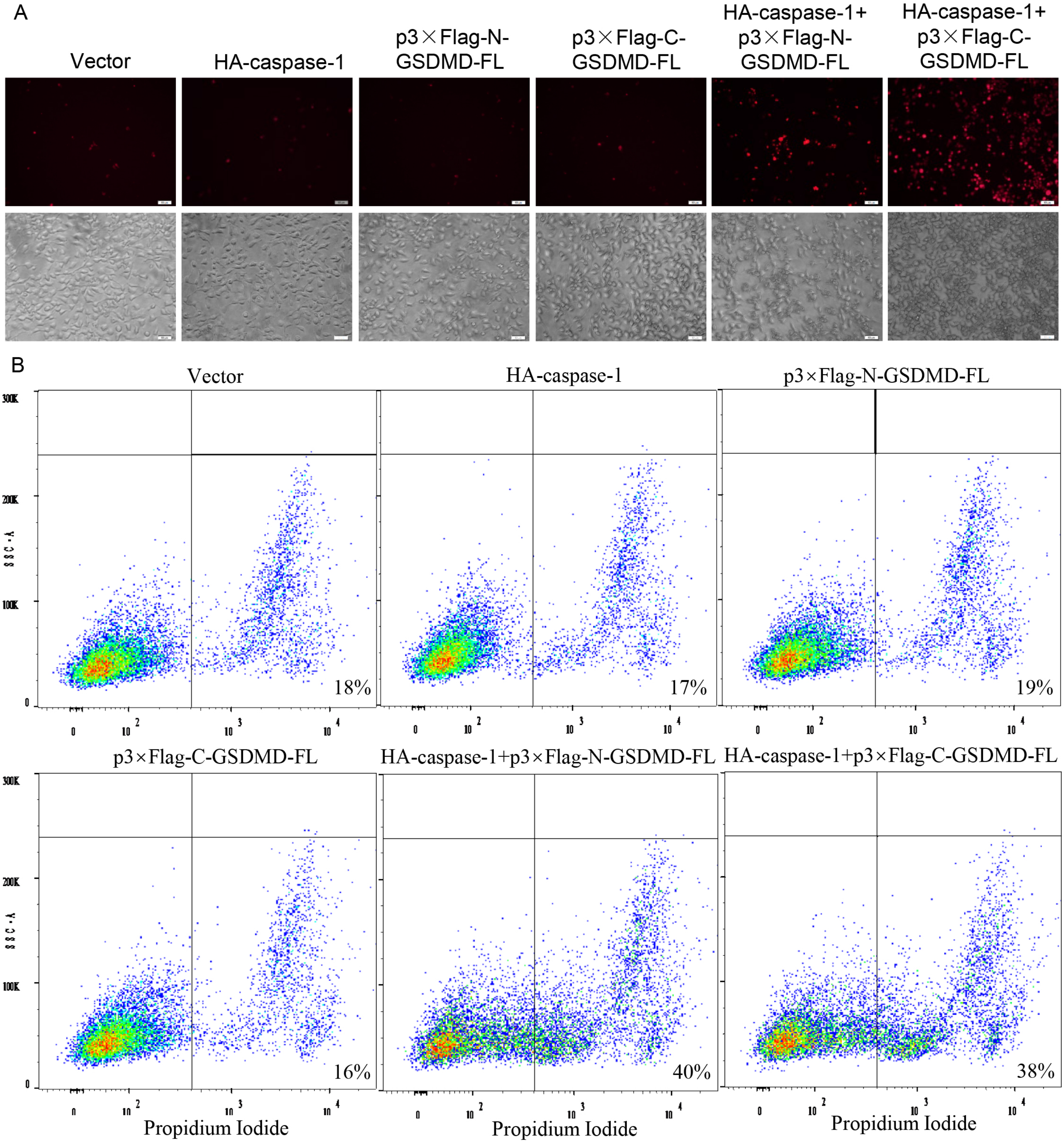

### Extended Data Figure 4

Extended Data Fig. 4

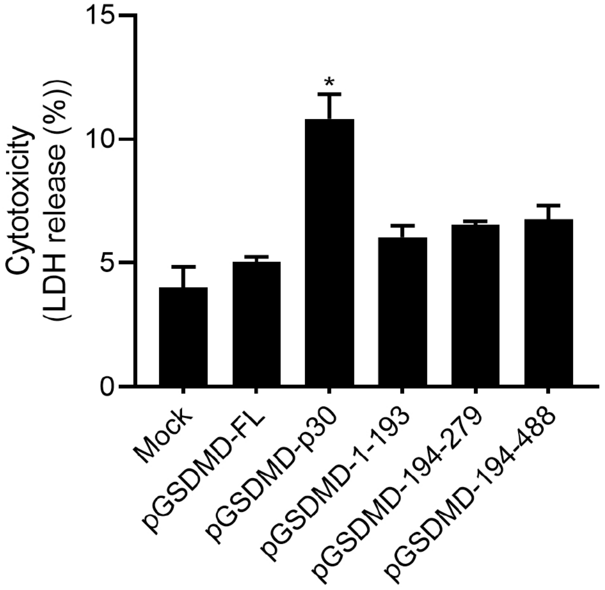


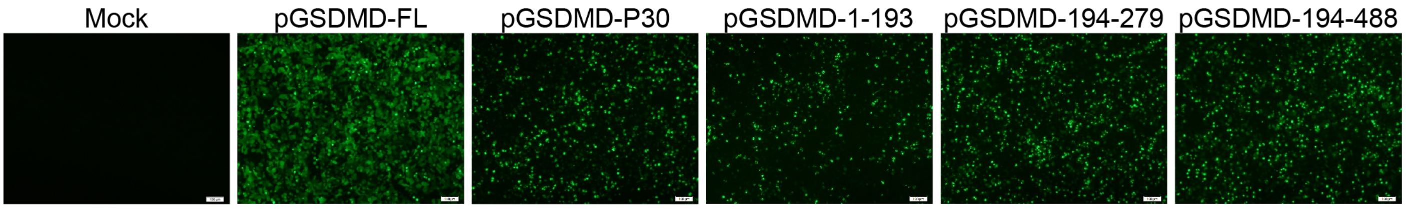

### Extended Data Figure 5

Extended Data Fig. 5

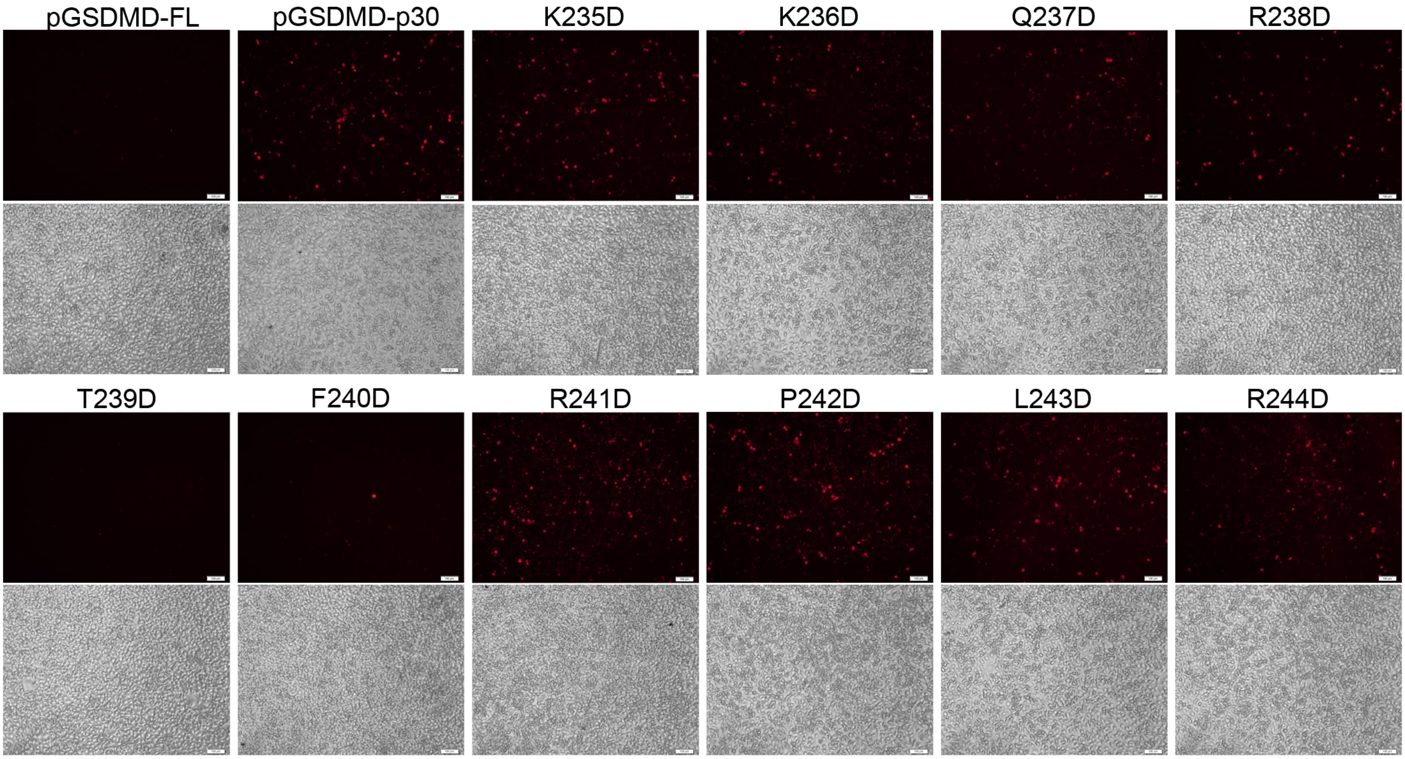

### Extended Data Figure 6

Extended Data Fig. 6

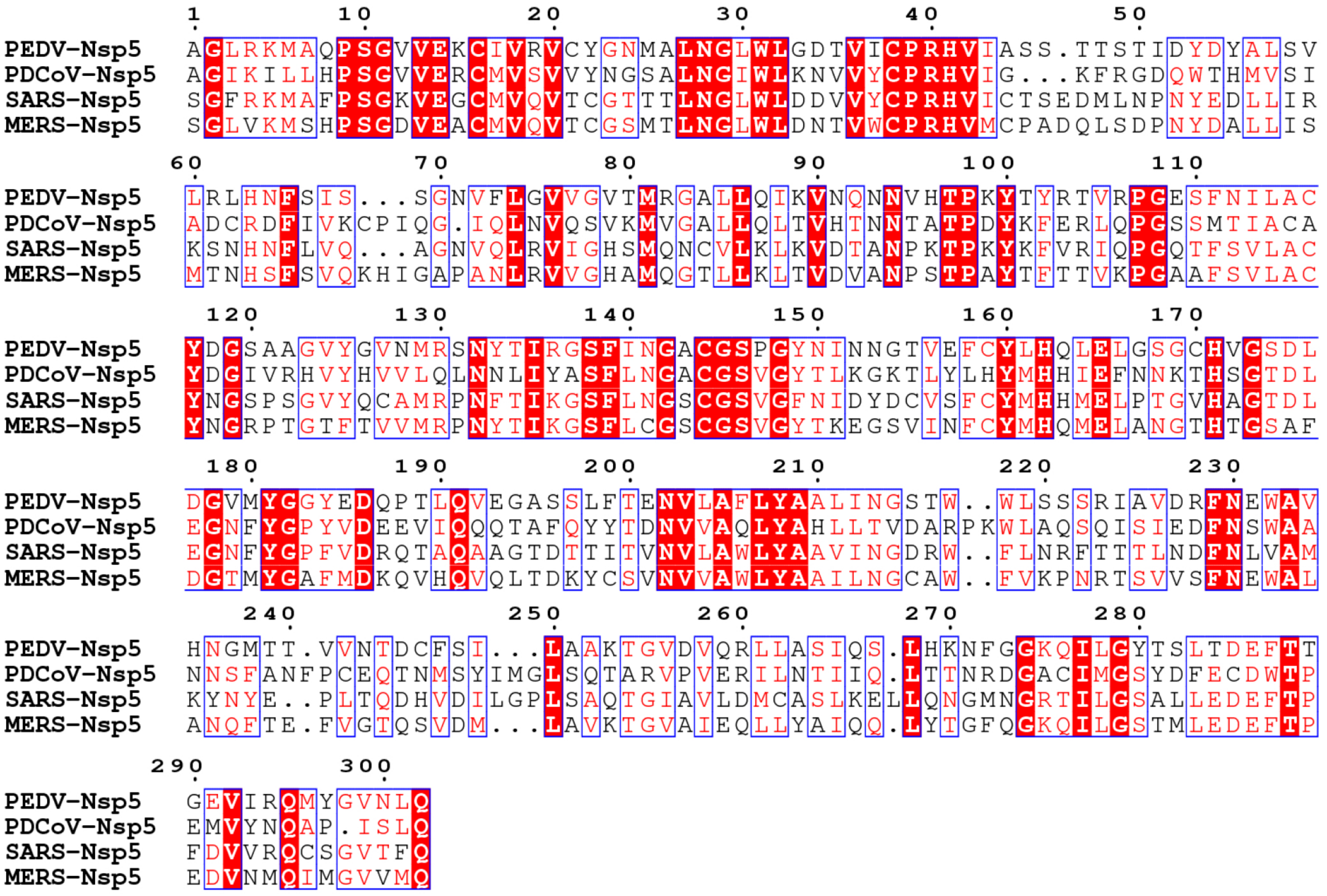
